# scFLUX: a web server for metabolic flux and variation prediction using transcriptomics data

**DOI:** 10.1101/2022.06.18.496660

**Authors:** Zixuan Zhang, Wennan Chang, Norah Alghamdi, Mengyuan Fei, Changlin Wan, Alex Lu, Yong Zang, Ying Xu, Wenzhuo Wu, Sha Cao, Yu Zhang, Chi Zhang

## Abstract

Quantitative assessment of single cell fluxome is critical for understanding the metabolic heterogeneity in diseases. Unfortunately, single cell fluxomics using laboratory approaches is currently infeasible, and none of the current flux estimation tools could achieve single cell resolution. In light of the natural associations between transcriptomic and metabolomic profiles, it remains both a feasible and urgent task to use the available single cell transcriptomics data for prediction of single cell fluxome. We present scFLUX here, which provides an online platform for prediction of metabolic fluxome and variations using transcriptomics data, on individual cell or sample level. This is in contrast to other flux estimation methods that are only able to model the fluxes for cells of pre-defined groups. The scFLUX webserver implements our in-house single cell flux estimation model, namely scFEA, which integrates a novel graph neural network architecture with a factor graph derived from the complex human metabolic network. To the best of our knowledge, scFLUX is the first and only web-based tool dedicated to predicting individual sample-/cell-metabolic fluxome and variations of metabolites using transcriptomics data. scFLUX is available at http://scflux.org/. The stand-alone tools for using scFLUX locally are available at https://github.com/changwn/scFEA.

## INTRODUCTION

Metabolic pathways provide essential energy and building blocks for the function of all cells, and dysregulated metabolism is a hallmark of many disease types such as cancer, diabetes, cardiovascular disease, and Alzheimer’s disease [1-7]. Given the pervasive role of metabolism in essentially every aspect of the disease pathology, an accurate and refined characterization of metabolic alterations and inference of their causes or downstream effects could have far-reaching impact in our knowledge on the basis of disease biology, clinical diagnosis and prevention, and disease management. Specifically, these include: (1) substantial increase of knowledge on metabolic variation, reprogramming and heterogeneity in the disease tissue microenvironment that accompany the disease initiation and progression [8, 9]; (2) identifying new drug targets or novel metabolic biomarkers for early diagnosis or therapeutic optimization [10, 11]; and (3) providing diet or nutrition recommendations for patients [12, 13].

Numerous computational methods have been proposed to study metabolic activities in different species [14-19]. However, while substantial efforts have been paid on reconstructing genome-wide metabolic maps, a fundamental question that remains un-addressed is how metabolic activities differ among cells of different morphological types, physiological states, tissues, or disease backgrounds that have the same genetic constitutions. Although transcriptomics or metabolomics experiments have been utilized to characterize metabolic alterations in diseases [20, 21], existing analysis tend to portray the average change of intermixed and heterogeneous cell subpopulations within a given tissue [22-24]. This makes it impossible to further study the metabolic heterogeneity and cell-wise flux changes in a complex tissue, in which cells are well understood to rewire their metabolism and energy production in response to varied biochemical conditions [25-28].

We have recently developed the first computational method to estimate cell-wise metabolic fluxome (flux distribution of the whole metabolic network) by using single-cell RNA-seq data (scRNA-seq) [29]. To the best of our knowledge, this is the first and only method that can robustly estimate flux distribution of a metabolic network at the resolution of individual cells. scFEA utilizes a factor graph base representation of metabolic network and a novel graph neural network (GNN) model for flux estimation, by assuming (1) metabolic flux can be modeled as a neural network of the genes involved in neighboring reactions, and (2) minimization of the flux imbalance of intermediate metabolites. Compared to existing methods, such as flux balance analysis or enrichment analysis-based approaches, scFEA is the only method that could (1) specifically model the nonlinear dependency between gene expression and metabolic flux, (2) assess flux in each single cell, and (3) allow for flux imbalance of certain intermediate metabolites in light of the high complexity of disease microenvironment.

Here we introduce scFLUX, a webserver that generalizes and implements the scFEA pipeline, in order to provide an optimized and coding-free environment to conduct single cell-or sample-wise metabolic flux estimation analysis by using transcriptomics data. Notably, scFLUX enables the input of both scRNA-seq and bulk RNA-Seq data to retrieve individual cell-or sample-wise metabolic flux prediction. In light of the complexity of the whole metabolic network, we manually curated the global metabolic map and a number of more focused metabolic subnetworks based on their topological structures for both human and mouse. Users can choose from the curated metabolic networks provided by the webserver. Existing webserver that has the closest scope of scFLUX is Fluxer [30]. However, Fluxer was designed for prokaryotes genomics data, focused on visualization and was based on Flux Balance Analysis model, which cannot predict sample-wise and disease context specific flux distribution, making it largely different from scFLUX. Within scFLUX, we also developed functionalities for downstream analysis of the predicted metabolic flux profiles. These include functions to compute: (i) levels of accumulation or depletion of metabolites and (ii) the subset of cells having distinct variation of certain metabolic modules [29].

## MATERIAL AND METHODS

### Reconstruction and representation of metabolic networks

The whole metabolic network in human and mouse have been well studied. However, while databases including the Kyoto Encyclopedia of Genes and Genomes (KEGG) [31] and Recon3D [32] provide well categorized metabolic pathways and the comprehensive set of metabolic genes, the network topological structure needs to be further optimized for fluxome estimation in single sample resolution by using transcriptomics data, due to the following reasons: (1) the flux balance relationships among different reactions could vary depending on the optimization goal or computational assumption, such as flux balance condition of carbon, redox or pH, (2) the network complexity needs to be reduced to enable computational feasibility, and (3) the low capture rate of mRNA abundance [33] of some metabolic genes may cause underestimations of flux in sample-wise analysis. In addition, cells of different types or physiological states naturally have varied metabolic characteristics.

In scFLUX, a metabolic network is first systematically decomposed and represented by connected **<u>metabolic modules</u>**, which is defined as a set of chain-shape reactions with (1) single input-/output-ends (reactions) and (2) no branch connected to other reactions from its intermediate reactions (**Fig 1**) [29]. The concept of metabolic module has been utilized by KEGG, in which many branches from intermediate metabolites were omitted [31]. In our reconstruction of metabolic modules in scFLUX, we (i) tightly followed the no-branch constraint, (ii) generalized the definition of a metabolic module by extending the single input- (or output-end) condition to be a class of metabolites which are not the output (or input) of other modules, i.e., at the boundary of the system, and (3) ensured a very small overlaps of enzymes shared by different modules [29]. Our new network reconstruction kept the major topological property of a metabolic network and optimized the factor graph for ease of computational prediction of metabolic fluxome using transcriptomics data (see details in Supplementary Information).

**Figure 1.**
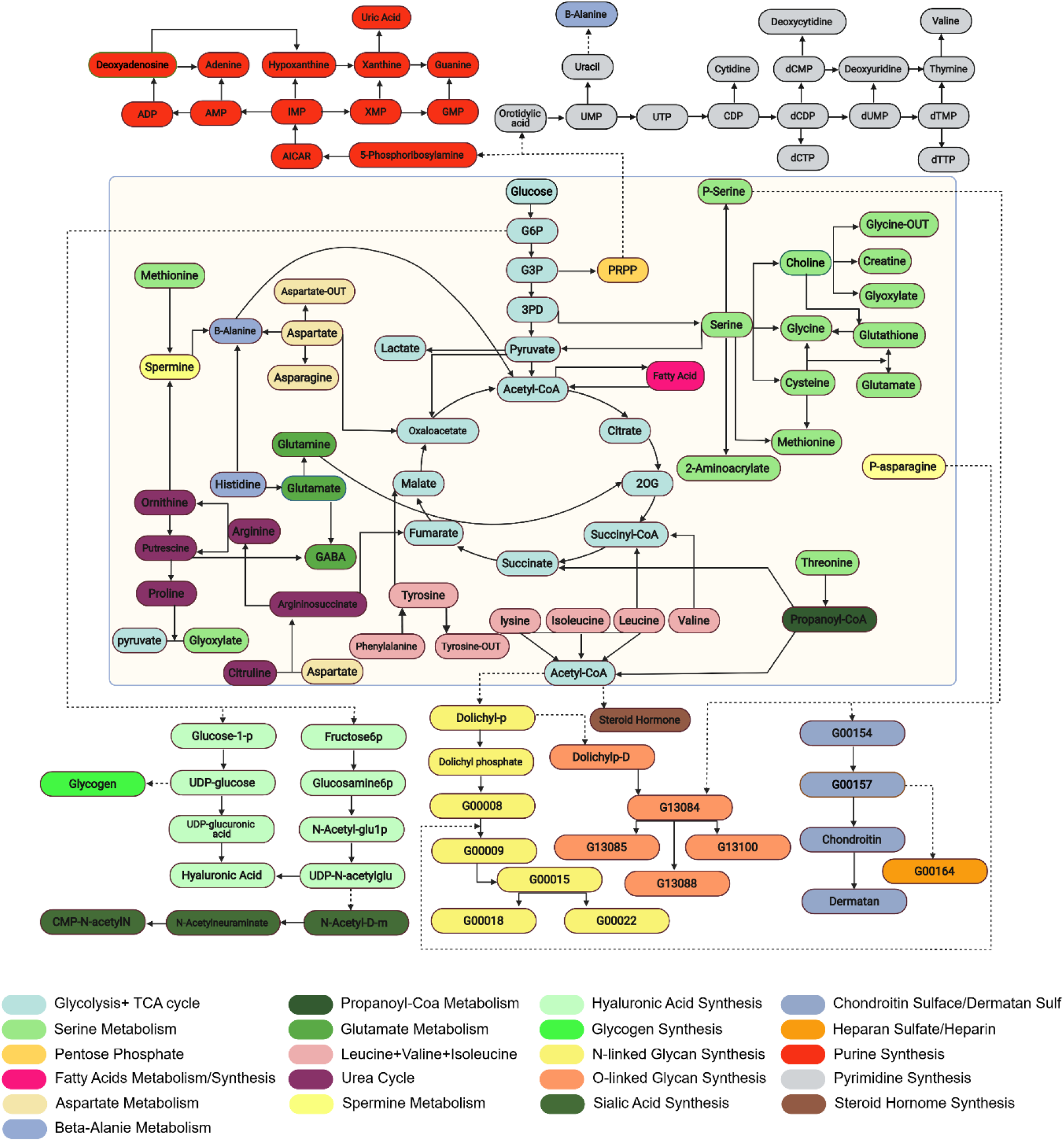
Reconstructed M171 metabolic map of human and mouse. Modules belong to different metabolic pathways (or termed super modules) are highlighted by different colors. Modules of metabolisms are presented in central yellow block while biosynthesis modules are presented outside. Glycans are labeled as KEGG G IDs.

Currently, scFLUX webserver houses the central metabolic map, namely M171, (an almost completed metabolic map), and eight smaller and more focused metabolic sub-networks, for both human and mouse (**Table 1**). Take the central metabolic map M171 as an example, which covers almost the complete metabolic network (**Figure** 1). We collected reactions of metabolism and biosynthesis as well as transporters for import and export from different data sources. Metabolic reactions were directly retrieved from KEGG [31]; the transporters and annotations of import and export reactions were accessed from the transporter classification database [34]; biosynthesis reactions were collected from the biosynthesis pathways encoded in KEGG and curated by using additional literatures.

**Table 1.** Reconstructed metabolic networks in scFLUX.

| Network Name | Network Description | #Modules | #Enzymes | #Intermediate Metabolites | #Genes (in Human, Mouse) |
| --- | --- | --- | --- | --- | --- |
| **M171** | An almost complete metabolic map | 171 | 451 | 70 | 663, 719 |
| **GlucoseGlutamineClose** | Glycolysis, TCA cycle, and glutaminolysis pathways | 23 | 98 | 17 | 132, 134 |
| **GlucoseGlutamineOpen** | General glucose and general glutamine metabolic pathways | 27 | 146 | 17 | 165, 176 |
| **BCAA** | Branched chain amino acids metabolic pathways | 14 | 52 | 6 | 60, 64 |
| **Acetylcholine** | Acetylcholine biosynthesis and metabolism | 15 | 30 | 6 | 80, 86 |
| **Dopamine** | Dopamine biosynthesis and metabolism | 9 | 30 | 5 | 23, 24 |
| **Histamine** | Histamine biosynthesis and metabolism | 6 | 18 | 3 | 23, 24 |
| **Serotonin** | Serotonin biosynthesis and metabolism | 8 | 18 | 4 | 24, 26 |
| **IronIon** | Sub-cellular localization specific metabolic network of iron ion | 15 | 27 | 8 | 141, 152 |

The final metabolic map covers the metabolism, transport, and biosynthesis of carbohydrates, amino acids, fatty acids and lipids, glycan, nucleic acids and other co-factors in human and mouse, including 663 human genes (719 mouse genes) of 451 enzymes and 116 transporters, 1,471 reactions, 1,561 metabolites. Eventually, the M171 network is simplified and reconstructed into a factor graph for the implementation of the flux estimation analysis, consisting of 171 modules of 22 super module classes that have 66 intermediate substrates (**Figure 1**). Here each super module is a manually curated group of modules of highly connected functions (See Supplementary Table S1). In addition to this whole metabolic map, we also manually curated 8 subnetworks covering (1) glucose and glutamine metabolism that provide a specific focus on energy metabolism, (2) branched chain amino acids metabolism, (3) metabolism of four types of neuron transmitters that support the analysis of central nervous systems, and (4) subcellular localization specific iron ion metabolism, which is the largest metabolic network of metal ion, in both human and mouse (**Table 1**). Detailed biological characteristics and statistics of the M171 and sub-networks are given in Supplementary Information and Supplementary Figures S1-7, and could also be found on the Download page of scFLUX.

Users can select the metabolic network on the Analysis page or download the detailed information of the networks on the Download page of scFLUX. As each module contains highly dependent metabolic reactions, they could be also used as analysis units similar to a pathway or gene set for enrichment-based analysis. We provide the gene information of each reconstructed module in .gmt format, which can be directly implemented with gene set or single-sample gene set enrichment analysis (GSEA or ssGSEA) [35, 36].

### Method overview of single cell-or sample-wise flux estimation analysis

scFLUX utilizes a novel graph neural network architecture as in scFEA to model cell-or sample-wise metabolic flux of each module by using their transcriptomic profiles [29]. A module-based network is formulated as a factor graph, where each module represents a factor, and each intermediate compound is a variable node carrying a likelihood function describing its flux balance. Denote *FG*(*C, R, E* = {*E*_*C*→*R*_, *E*_*R*→*C*_}) as the factor graph, where *C* is the set of metabolites, *R* is the set of metabolic modules, *E*_*C*→*R*_ and *E*_*R*→*C*_ represent direct edges from module to metabolite and from metabolite to module, respectively. To illustrate the flux estimation method, we use the “GlucoseGlutamineOpen” network encoded in scFLUX as an example, which contains 27 modules and 17 intermediate metabolites that covering the glycolysis, TCA cycle and glutamine metabolic pathways (**Figure 2**). For each intermediate metabolite *C*_*k*_ in this network, define the set of modules consuming and producing each *C*_*k*_ 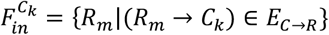 and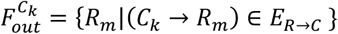. For example, 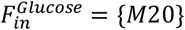 and 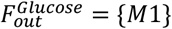 in this network. For a transcriptomics dataset containing *N* samples, denote 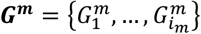 as the genes involve in the module *R*_*m*_, 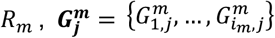 as their expression and *Flux*_*m,j*_ as the flux of the module *m* in the cell or sample *j*. We 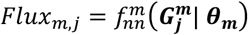 as a multi-layer fully connected neural network with the input 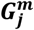, where ***θ***_***m***_ denotes the parameters of the neural network (**Figure 2**). Then the *θ*_***m***_ and cell-or sample-wise flux *Flux*_*m,j*_ are solved by minimizing the following loss function:

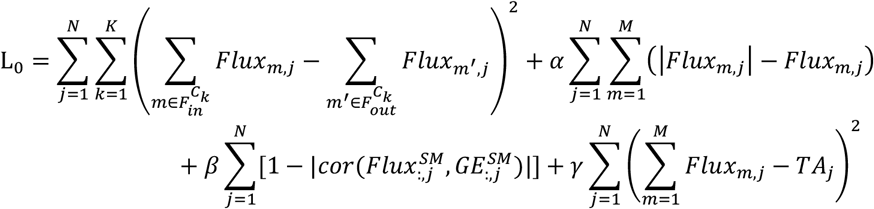

, where *α, β*, and *γ* are hyperparameters, *cor* represents Pearson correlation coefficients; *Flux*^*SM*^ and *GE*^*SM*^ are two *NSM* × *N* matrices, here *NSM* is the number of super modules, 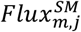 represents the sum of the flux of the modules in the super module *m*, 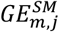 represents the sum of expression of the genes in the super module *m*, in cell or sample *j*, and *TA*_*j*_ is a surrogate for total metabolic activity level of cell or sample *j*, which is assigned as the total expression of metabolic genes in cell or sample *j*. Hence, the first, second, third and fourth terms of *L* correspond to constraints on flux balance, non-negative flux, the coherence between predicted flux and total gene expression level of each super-module, and the relative scale of flux, respectively.

**Figure 2.**
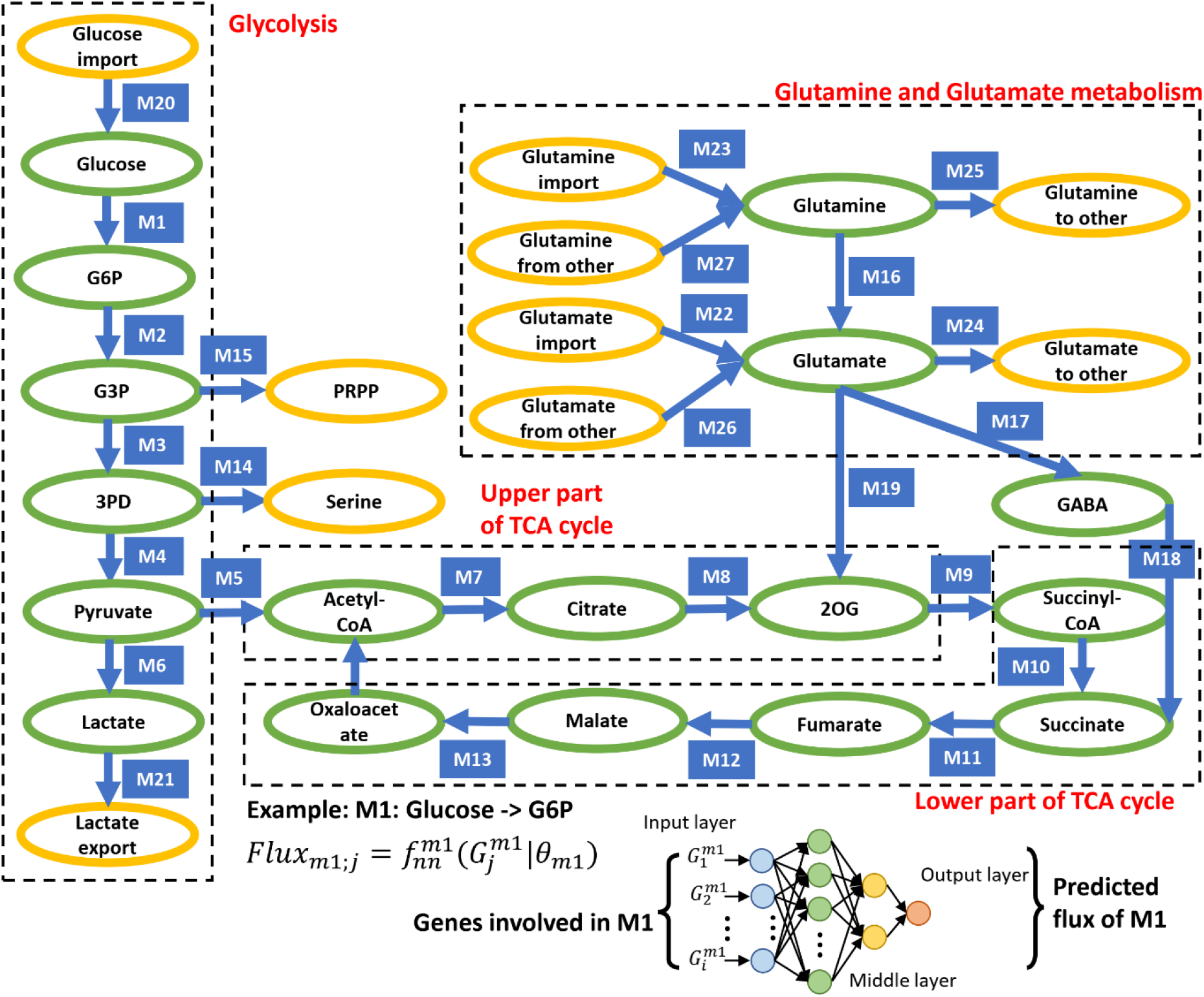
Factor graph representation of the subnetwork of Glycolysis + TCA cycle + Glutamine metabolism. Modules (variables) and Metabolites (factors) are represented by rectangles and ovals, respectively. Intermediate and end metabolites are green and yellow colored, respectively. An example of the flux model of M1: glucose -> G6P is given in the bottom, which illustrates how the flux of a module depends on the gene expression involved in the module.

The above flux estimation model has been validated on two sets of matched scRNA-seq and bulk cell metabolomics data, simulated scRNA-seq and fluxome data, and 5 sets of high quality tissue transcriptomics or scRNA-seq datasets [29]. We have tested the robustness of scFEA regarding its hyperparameters with other users on more than 20 scRNA-seq datasets. Our robustness analyses suggested that an empirical setting of *α* = 1, *β* = 0.1, *γ* = 1 can guarantee good prediction accuracies of reasonable biological interpretability, with fast convergence rate (see detailed discussion in Supplementary Information).

### scFLUX web server

scFLUX is user-friendly web server that provides a coding free environment for users to conduct end-to-end single cell-or sample-wise flux estimation analysis using transcriptomics data. As illustrated in **Figure 3**, scFLUX takes as inputs a single cell or bulk tissue transcriptomics data, and a user defined metabolic network, conducts data preprocessing and flux estimation steps, and outputs cell-or sample-wise metabolic flux estimations and their variations, as well as result annotation files. The web server implementation is described below, with detailed information provided in Supplementary Information.

**Figure 3.**
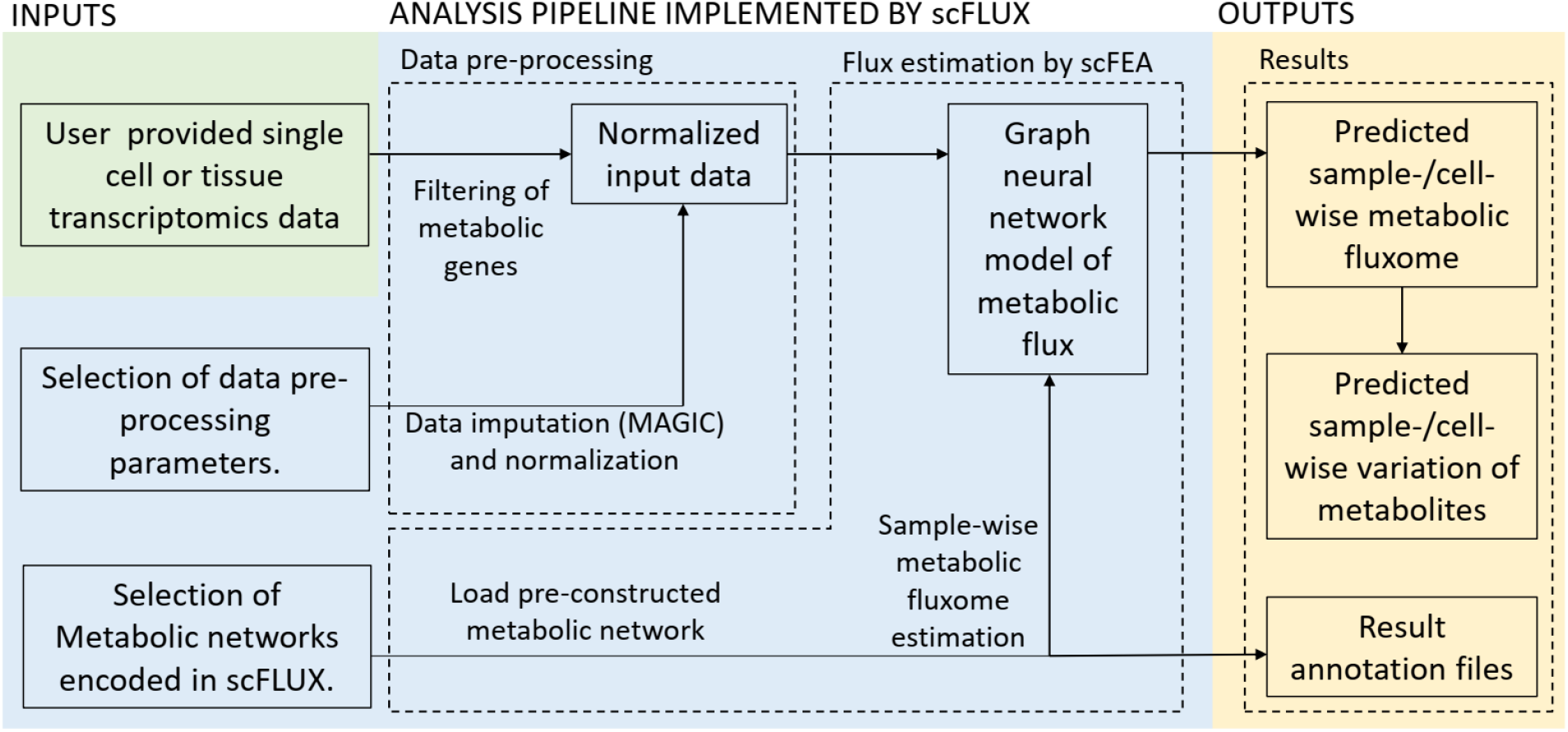
Flowchart of the scFLUX webserver. The inputs, analysis pipeline and outputs are listed in green, blue, yellow blocks, respectively.

*Front-end*. The scFLUX web-based server is implemented in Python by using the Django framework. scFLUX utilizes the SQLite database for a persistent storage and retrieval of requested input or output data. The Nginx HTTP server is utilized as a secure application gateway, which optimizes data uploading, serves as a reverse proxy, and provides a caching mechanism. The python program of scFLUX is deployed via the uWSGI server and the program communicates with uWSGI by the WSGI spec. Here the uWSGI server enables an efficient management and resource allocation for multiple processes. The website interface was designed by using the Bootstrap framework, the jQuery JavaScript library, and extension packages.

*Back-end*. Data input and processing are conducted by using Pandas, NumPy, and Pyreadr. Data preprocessing procedure consists of three major steps: (1) an evaluation step that checks if the input file is in a correct format. Warnings will be returned if the data format does not meet the requirement of scFLUX; (2) data normalization and imputation by using MAGIC [37]; and (3) passing the processed expression data and user selected metabolic network to the flux estimator.

The flux estimation procedure is conducted by using PyTorch, Pandas and NumPy. The factor graph model of each metabolic network is first built, in which each module is a variable, and each intermediate metabolite is a factor. Parallel three-layer neural networks were first constructed to model the non-linear dependency between gene expression involved in the modules and their flux rates. Noted, one neural network is constructed for each module. The loss function is further constructed based on the in-/out-flux of each intermediate metabolites and minimized by using the Adam method, which is the most efficient stochastic optimization approach in the built-in optimizer of Pytorch. The loss curves are drawn by using Matplotlib and shown on the Results page (**Figure 4**).

**Figure 4.**
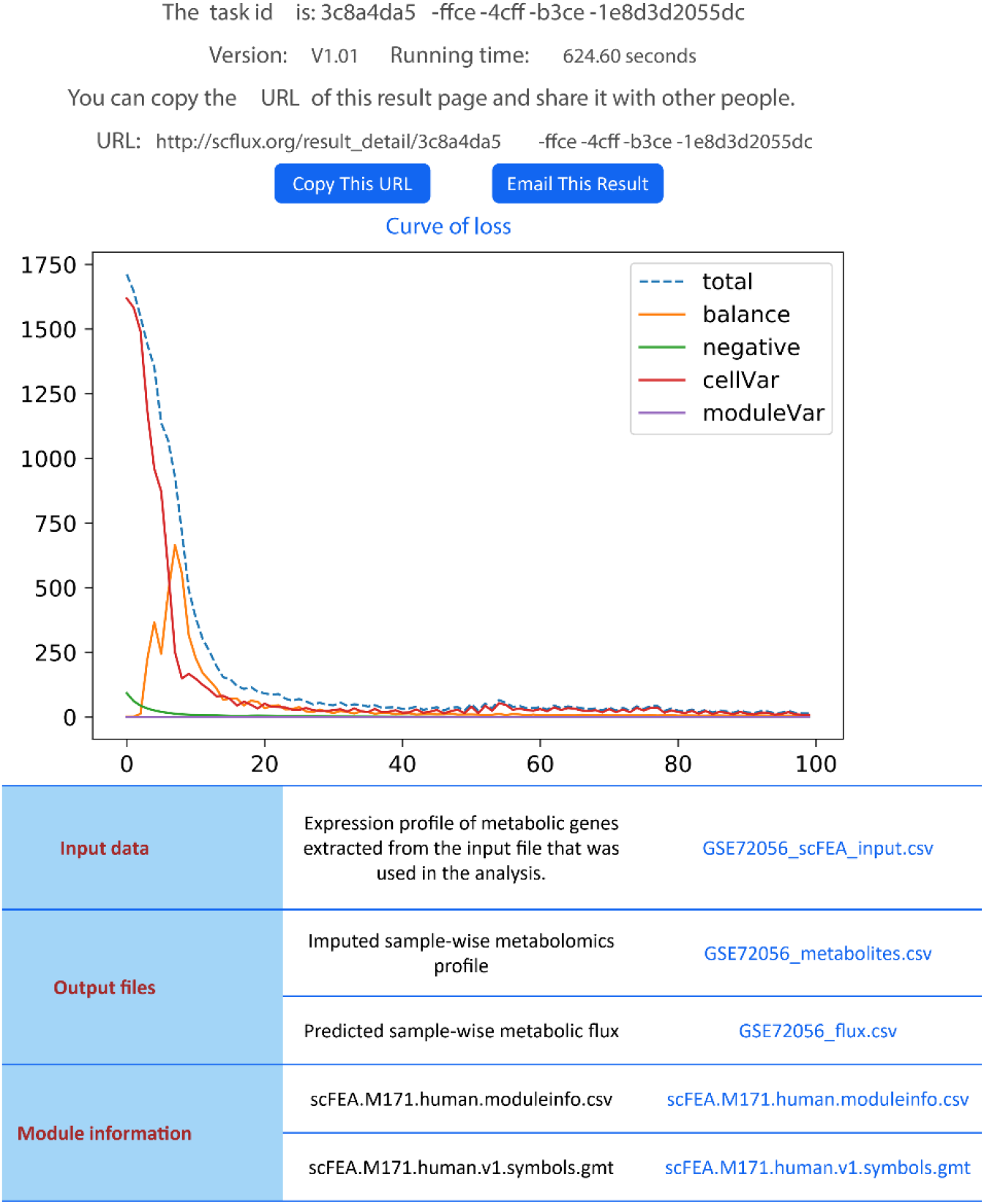
Results page of scFLUX.

*Job Management*. scFLUX manages asynchronous tasks by using Celery via a distributed task queue. Users can run multiple analysis tasks simultaneously. A unique task ID and link will be generated when a task is submitted, by which the user can track the progress of the analysis task. When the task is completed, the user can use the link to access, download or share the analysis results. Anyone with the task ID (or link) can access the results. Analysis results will be stored on the server for at least one year. The list of user’s tasks is implemented by using a cookie mechanism with a consent form provided on the web page. If the cookie is enabled by a user, the Results page will list previous tasks submitted by this user on the same browser.

*Browser compatibility*. scFLUX has been tested on major modern browsers including Google Chrome, Mozilla Firefox, Safari and Microsoft Edge.

## RESULTS

### Server input

The input of scFLUX includes (1) a transcriptomics data set containing at least 25 samples and (2) a species-specific metabolic network and analysis parameters selected by users. The scFLUX webserver is currently housing the central metabolic map (M171) and eight specific metabolic sub-networks of human and mouse. Users could select among the two species and the nine metabolic networks from the boxes on the left-hand side of the home page. The input scRNA-seq or general RNA-Seq dataset should have genes on its rows, and samples on its columns, and TPM (or CPM/FPKM) normalized data is recommended. scFLUX webserver accepts comma-(.csv), space-(.txt), or tab-(.txt) delimited input files, which should be in a matrix format and contain row/column names. For row names, both gene symbol and Ensembl gene ID of human and mouse are accepted. The maximal input file size is 500MB. For a large data set, we recommend users upload only the gene expression data of the scFLUX metabolic genes, as other genes will not be used in the flux estimation. The metabolic genes used for flux estimation in scFLUX can be downloaded from the Download page for both human and mouse. Notably, the graph neural network-based formulation of scFLUX allows for the flexibility that expression values of some metabolic genes may be missing from the input data.

scFLUX provides two pre-processing options for the input transcriptomics dataset: (1) Imputation. If the input transcriptomics data, such as an scRNA-seq data, is highly sparse, an imputation procedure is recommended. The default imputation method is MAGIC [37]. (2) Normalization. Four options are provided by scFLUX: (i) no normalization, (ii) log transformation, namely log(x+1), where x represents the original input expression matrix, (iii) CPM normalization, and (iv) log transformation of CPM normalized values, namely, log(CPM+1). Users need to decide whether and how these two procedures will be performed by specifying two relevant hyperparameters on the running page of scFLUX.

The input transcriptomics data will be checked by a format validator, and a job will be submitted only when the input file meets the requirements of scFLUX. To help users submit their tasks correctly, scFLUX provides sample input files on the home page.

### Server output

When an analysis task is completed, the user can obtain the output files of the job on its Results page (**Figure 4**). For each task, scFLUX provides four downloadable results as follows:

1. The predicted cell-or sample-wise metabolic flux. It is a .csv file with modules on its row, and single cells or samples on its column, and each entry is predicted flux rate of the metabolic module in the corresponding cell or sample.
2. The imputed cell-or sample-wise metabolomics profile. It is a .csv file with metabolites on its row, and samples on its column, and each entry is an imputed metabolomics change of the metabolite in the corresponding cell or sample.
3. Module information. It is a .csv file where each row contains the detailed information of the reactions and metabolic compounds involved in each module. These modules are the ones involved in the user defined metabolic network.
4. Module-gene information. It is a .gmt file where each row contains the gene symbols of the genes in each module of the analysed metabolic network. These genes are the ones used in the flux estimation process.

The users can view or download the convergence curves of the optimization process, for both the total loss, as well as the four individual loss terms in the optimization function. The predicted cell-or sample-wise metabolic flux is amenable to myriad downstream analyses, for which a number of functions are provided by the scFLUX website under Tutorial/Package tutorial section. Basically, the estimated flux matrix could be seamlessly integrated into the standard Seurat analysis pipeline for comparative analysis, dimensional reduction, clustering, as well as visualization. In addition, functions are supplied by scFLUX to assess levels of accumulation or depletion of metabolites, and detect subsets of cells or samples having distinct variation of certain metabolic modules.

### Case study

We have previously validated the core method of scFLUX, namely scFEA, using matched scRNA-seq and metabolomics data generated in house and public data. We have demonstrated that scFEA can accurately predict cell-or sample-wise flux and metabolomics changes, which largely outperforms pathway-enrichment based methods for metabolic pathway activity prediction [29] (see Supplementary Information). As no additional matched scRNA-seq and metabolomics data is available for a method validation, here we demonstrate the utility of scFLUX on (1) a collection of scRNA-seq datasets collected from human and mouse cancer microenvironment and (2) ROSMAP single nuclei RNA-seq (snRNA-seq) data collected from brain tissues of Alzheimer’s disease (AD) patients and healthy donors [38].

*Cancer data*. We applied scFLUX to scRNA-seq data collected from cancer microenvironments that contains cancer, myeloid and T cells, including eight human and two orthotopic mouse data sets, for flux estimation of the GlucoseGlutamineOpen network. Our analysis identified that: (i) cancer cells consistently have the highest glucose metabolic rate, including lactate production, TCA cycle, and nucleotide and serine biosynthesis, followed by myeloid cell and then T cells, in most human cancer and injected mouse tumor tissues analyzed (Supplementary Figure S8A,B); (ii) the rates of the total glucose consumption strongly correlate with the rates of proliferation (Supplementary Figure S8C); and (iii) the glycolytic flux distributions in human cancer and injected mouse tumor cells are considerably different, particularly in terms of the fractions into nucleotide and serine biosynthesis, which matches existing knowledge in (1) the roles played by serine in transplant rejection [9] and (2) the prevalently increased nucleotide biosynthesis in proliferating cancer cells [10].

*ROSMAP AD data*. We also applied scFLUX on ROSMAP snRNA-seq data to predict AD-specific metabolic variations by using the M171 network. We have identified that metabolic activity is higher in neuron cells than in other brain cell types. We further focused on the metabolomic changes predicted by scFLUX. Supplementary Table S2 lists 14 metabolites whose concentrations have the largest distinctions for neuron cells from AD and healthy control brain. Among them, increased glycolytic substrates and GABA, and decreased glucose, nucleic acids and branched chain amino acids (valine, leucine and isoleucine) have been reported [39, 40], while aspartate, serine and methionine may serve as new biomarkers [29].

## DISCUSSION

In this paper, we present scFLUX, the first and only web server for metabolic flux estimation for single cells or samples using single cell or bulk RNA-Seq data. The key methodology behind this server is based on our previous works on factor graph representation of a complex metabolic network, as well as a graph neural network-based solver for single cell fluxome estimation [29]. Noted, scFLUX conducts an end-to-end prediction and provides a ready means of interrogating the flux rate of metabolic modules and concentration changes of metabolites, which can be directly utilized to understand possible metabolic reprogramming events and guide targeted metabolomics experiments. We anticipate the applications of scFLUX could increase our understanding in (1) key metabolic reprogramming events and causes, and (2) the impact of metabolic abnormalities to other biological characteristics, which together will contribute to improved precision medicine regime such as biomarker screening and drug target prediction.

In order to best characterize context specific metabolic activities, there is still a few unsolved challenges. A complex tissue microenvironment may be constituted by cells of different metabolic abnormalities, heterogeneous metabolic networks, varied preferences, and dependencies [41-45]. In future work, we will enable the reconstruction of context specific metabolic networks and modules, especially disease, tissue and cell type specific ones, to maximize the discoveries of hidden and dynamic relationships among the metabolic units under different biological conditions. In addition, recent evidence suggests that the direction of certain reversible reactions may not be constant for cells within one disease microenvironment, which represents one way for the cells to increase their fitness level by reprogramming the metabolic exchange mechanisms under a highly perturbed environment [46]. Hence, a second future direction is to enable the assessment of sample-wise directions of reversible reactions and inter-cell metabolic exchange or competition by using single cell data. We will also extend flux estimation capability for other omics data types, like proteomics or metabolomics data. A few newly developed analysis features, including a perturbation analysis to determine the contribution of each gene to each flux, and newly curated metabolic modules including methionine and copper ion metabolic pathways, are currently available in the stand-alone version of scFLUX. Such features will be updated to scFLUX web server after thorough validations were conducted.

## AVAILABILITY

scFLUX is available as a web server at http://scflux.org/. The stand-alone tool package to run scFLUX on a local machine is available at the GitHub repository (https://github.com/changwn/scFEA).

## SUPPLEMENTARY DATA

Supplementary Data are available at NAR online.

## ACKNOWLEDGEMENTS

C.Z and S.C want to thank Dr. Yunlong Liu, Dr. Kun Huang and Dr. Xiongbin Lu for their constructive suggestions and advice to this work.

