## Supplementary material for "scFLUX: a web server for metabolic flux and variation prediction using transcriptomics data": Supp File

### Module and factor graph base representation of metabolic networks

In scFLUX, a metabolic network is first systematically decomposed and represented by connected **metabolic modules**, which is utilized by KEGG and defined by a set of chain-shape reactions with (1) single input-/output-ends (reactions) and (2) no branch connected to other reactions from its intermediate reactions (as illustrate in the figure on the right). In scFEA, we generalized the definition of a metabolic module by enabling the single input- (or output-end) be a class of metabolites if these metabolites are not the output (or input) of other modules, i.e., at the boundary of the system. The rationale of the generalized definition of output is that if the input (or output) end of a module does not link with any output (or input) of other module, no flux balance constraint will be set for it input (or output) end. Hence, the module serves as the boundary of the system, and its flux just reflect the total flux from any possible input (or output) to its output (or input), which can be estimated by the self-constrained model utilized in scFLUX.

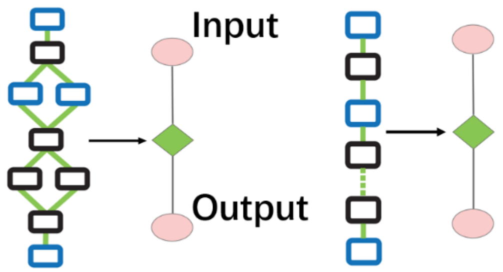

**An illustration of chain-shape modules**

### Reconstruction of metabolic network

*M171 Genome-wide metabolic network*

The M171 metabolic map consists of pathways and reactions that fall under four major types, namely import, metabolism, biosynthesis, and export in human and mouse. To ensure a comprehensive coverage of the global metabolic map, we collected reactions of metabolism and biosynthesis as well as transporters for import and export from different data sources. Specifically, metabolic reactions were directly retrieved from KEGG database [1]; the transporters and annotations of import and export reactions were accessed from the transporter classification database [2]; biosynthesis reactions were collected from the biosynthesis pathways encoded in KEGG and curated by using additional literatures (see details in Supplementary Methods). The final central metabolic map covers the metabolism, transport, and biosynthesis of carbohydrate, amino acids, fatty acids and lipids, glycan, and nucleic acids in human and mouse, including 663 human genes (719 mouse genes) of 451 enzymes and 116 transporters, 1471 reactions, 1561 metabolites. Noted, we did not split fatty acids metabolism and biosynthesis pathways into sub modules as they have a strong overlap of enzymes. We divided central metabolic into 22 super modules which including the glycolysis and TCA cycle super module, the serine metabolism super module, the pentose phosphate super module, the fatty acids metabolism super module, the aspartate metabolism super module, the beta-alanine metabolism super module, the propionyl-CoA metabolism super module contains, the glutamate metabolism super module, the leucine, valine and isoleucine super module contains, the urea cycle super module, the spermine metabolism super module, the transporters super module, the glycogen synthesis super module, the glycosaminoglycan synthesis super module, the N-linked glycan synthesis super module, the O-linked glycan synthesis super module, the sialic acid synthesis super module, the purine synthesis super module, the pyrimidine synthesis super module, the steroid hormone synthesis super module. The detailed statistics of the M171 network shows in **Supplementary Table S1**.

| Network Name | Network Description | #Modules | #Enzymes | #Intermediate Metabolites | #Genes (Human, Mouse) | #Reactions | #Total Metabolites |
|---|---|---|---|---|---|---|---|
| **M171** | An almost complete metabolic map | 168 | 451 | 70 | 663, 719 | 1471 | 1561 |
| **GlucoseGlutamineClose** | Glycolysis, TCA cycle, and glutaminolysis pathways | 27 | 146 | 17 | 176, 165 | 344 | 441 |
| **GlucoseGlutamineOpen** | General glucose and all glutamine metabolic pathways | 23 | 98 | 17 | 132, 134 | 243 | 292 |
| **BCAA** | Branched chain amino acids metabolic pathways | 14 | 52 | 6 | 60, 64 | 207 | 261 |
| **Acetylcholine** | Acetylcholine biosynthesis and metabolism | 15 | 30 | 6 | 80, 86 | 79 | 109 |
| **Dopamine** | Dopamine biosynthesis and metabolism | 9 | 30 | 5 | 23, 24 | 132 | 193 |
| **Histamine** | Histamine biosynthesis and metabolism | 6 | 18 | 3 | 23, 24 | 88 | 135 |
| **Serotonin** | Serotonin biosynthesis and metabolism | 8 | 18 | 4 | 24, 26 | 156 | 243 |
| **IronIon** | Sub-cellular specific metabolic network of iron ion | 15 | 27 | 8 | 141, 152 | 51 | 114 |

**Supplementary Table S1. Detailed statistics of M171 network.**

In addition to the M171 genome-wide metabolic network, we also manually curated 8 subnetworks of (1) glucose and glutamine metabolism that provide a specific focus on energy metabolism, (2) branched chain amino acids metabolism, (3) metabolism of four types of neuron transmitters that support the analysis of central nervous systems, and (4) subcellular localization specific iron ion metabolism, which is the largest metabolic network of metal ion, in both human and mouse, as detailed below. Detailed network information can be downloaded via http://scflux.org/.

*Central Energy Metabolic Network*

The GlucoseGlutamineClose (**Supplementary Figure S1**) and GlucoseGLutamineOpen (**Figure 2** of the main text) networks focus on central energy metabolic pathways that covers glycolysis and other metabolism of glucose, TCA cycle and glutamine metabolism, which are differed by assumptions on glutamine and glutamate metabolism. Specifically, the GlucoseGlutamineClose models assume that glutaminolysis pathway is the main branch that consumes glutamine and glutamate in the system and the influx of glutamine and glutamate majorly replies on import by transporters, which is designed to evaluate the relative ratio of glycolysis and glutaminolysis. On the other hand, GlucoseGlutamineOpen network assumes there are additional sources that exchange glutamine and glutamate, which characterize the comprehensive glutamine and glutamate metabolism.

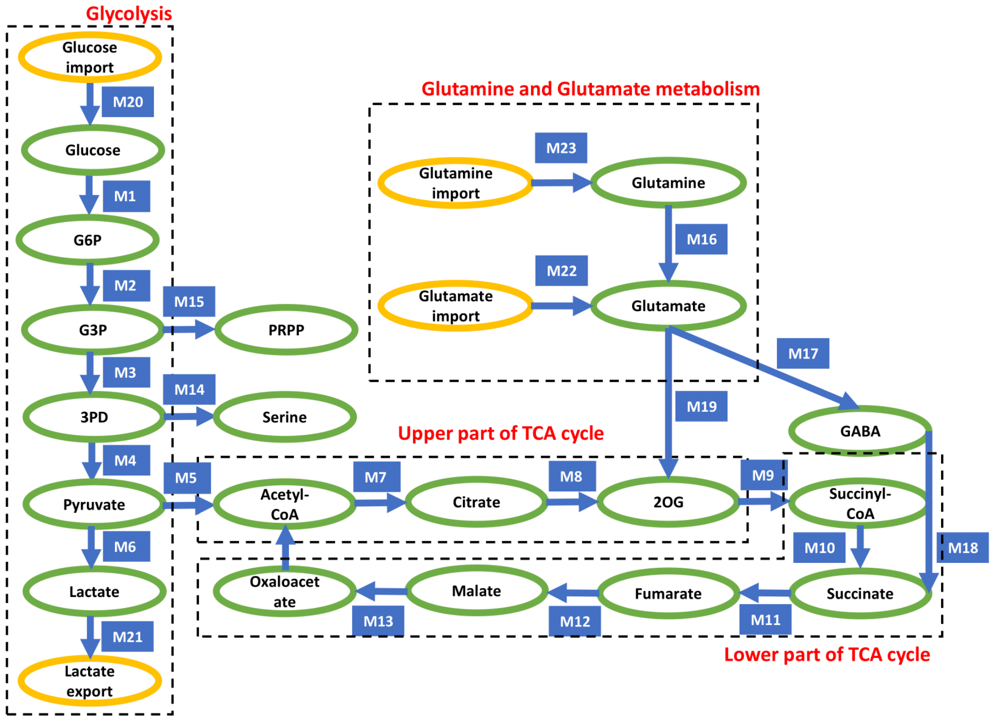


**Supplementary Figure S1. The GlucoseGlutamineClose Network.**

*Branched Chain Amino Acids metabolism*

The BCAA network is of the network of Branched Chain Amino Acids (BCAA) metabolism (**Supplementary Figure S2**). We have collected metabolic reactions of leucine, valine and isoleucine metabolism, biosynthesis of branched chain fatty acids, and downstream metabolism to acetyl-CoA and propanoyl-CoA.

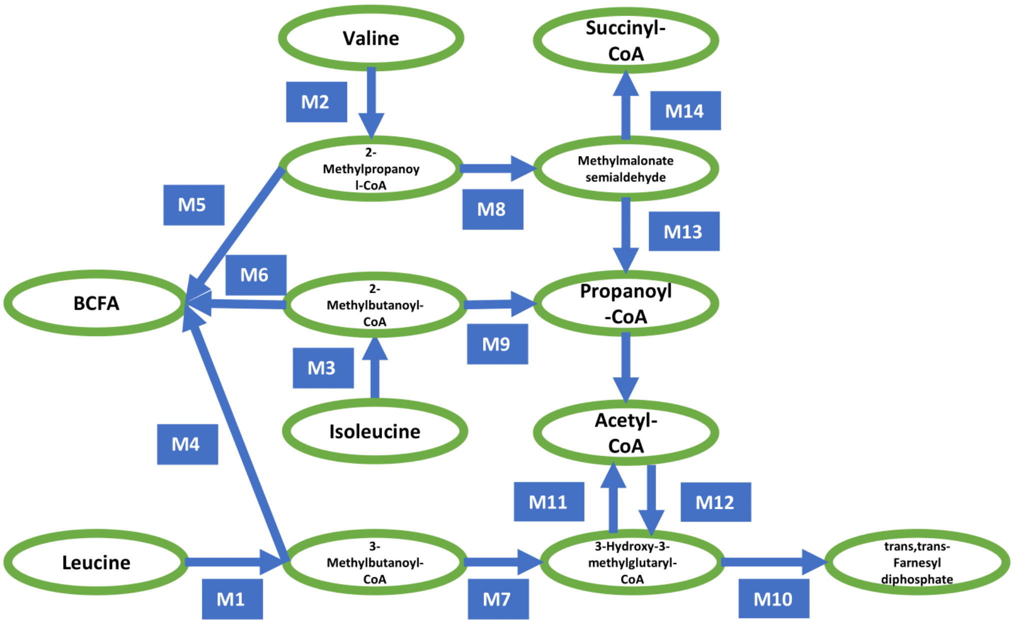


**Supplementary Figure S2. BCAA metabolic Network.**

*Neuron transmitter metabolism*
The Acetylcholine network is the network of biosynthesis and metabolism of the neurotransmitter acetylcholine (**Supplementary Figure S3**). We have collected related reactions from glycerol 3-phosphate to acetylcholine and its metabolism into choline.

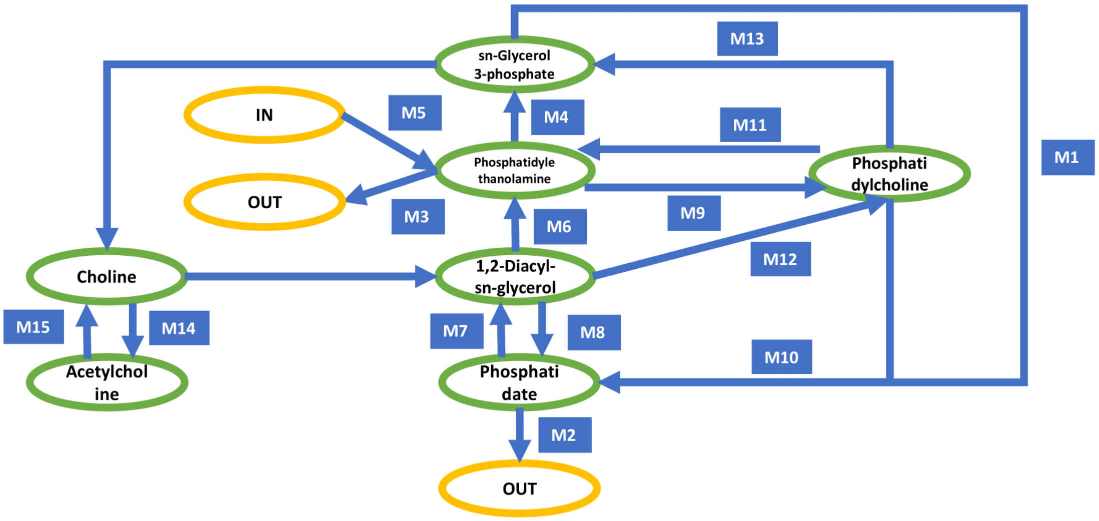


**Supplementary Figure S3. Acetylcholine metabolic Network.**

The Dopamine network is the network of biosynthesis and metabolism of the neurotransmitter dopamine (**Supplementary Figure S4**). We have collected related reactions from tyrosine to dopamine and its metabolism/biosynthesis into downstream products such as noradrenaline and adrenaline.

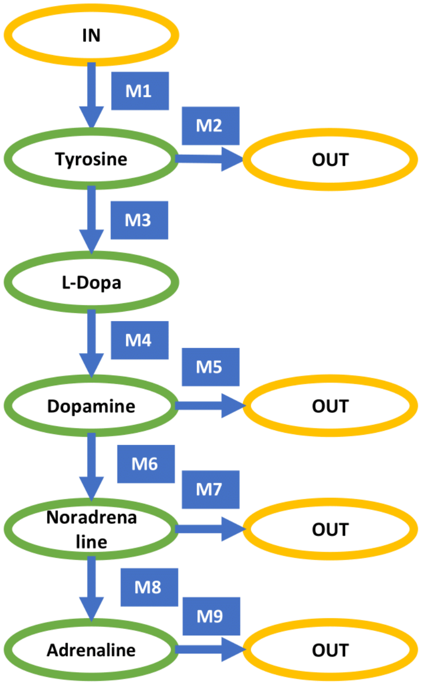


**Supplementary Figure S4. Dopamine metabolic Network.**

The Histamine network is the network of biosynthesis and metabolism of the neurotransmitter histamine (**Supplementary Figure S5**), which also serves as an organic nitrogenous compound involved in local immune responses. We have collected related reactions from histidine to carnosine and histamine.

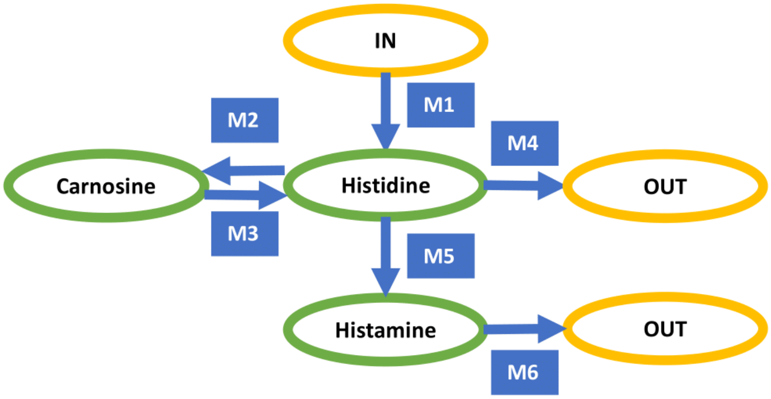


**Supplementary Figure S5. Histamine metabolic Network.**

The Serotonin network is the network of biosynthesis and metabolism of the neurotransmitter serotonin (**Supplementary Figure S6**). We have collected related reactions from tryptophan to oxitriptan and serotonin, and its downstream metabolism into melatonin and degradation.

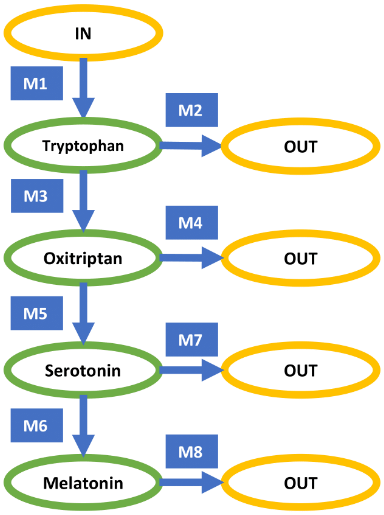


**Supplementary Figure S6. Serotonin metabolic Network.**

The IronIon network is a sub-cellular specific metabolic network of iron ion (**Supplementary Figure S7**). We have collected the reactions including iron ion transportation, ferric ion reduction, and utilization of iron ion in heme and ion sulfur biosynthesis and Fenton reaction. The constructed map of iron metabolic reactions in a human cell consists of three sources to the cytosolic $Fe^{2+}$ pool, namely ferrous ion import, ferric ion import, and reduction, and heme import and reduction; four sinks for the cytosolic $Fe^{2+}$, namely mitochondrial Fe-S cluster and heme synthesis, ferrous ion export, and Fenton reaction; and sources and sinks of the $O_2^{\cdot -}$ and $H_2O_2$, totaling eight. The fifteen metabolic branches were each considered as a metabolic module, each containing one to a few dozen of metabolic genes, whose expression levels were utilized to estimate their metabolic flux.

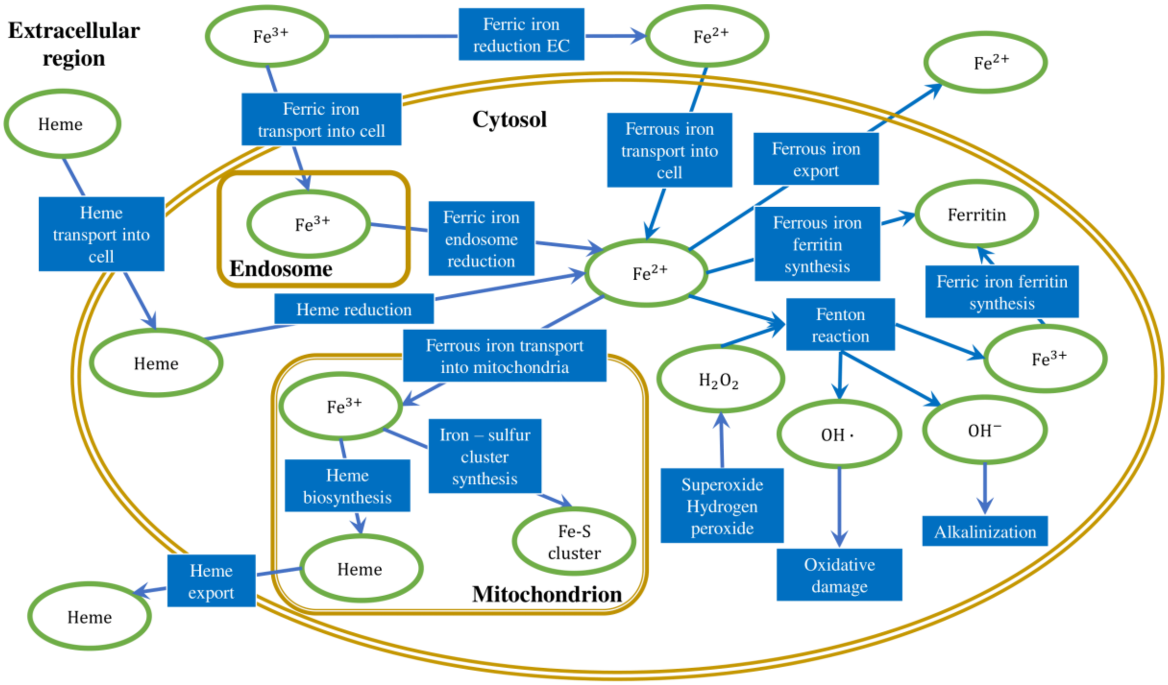


**Supplementary Figure S7. Subcellular localization specific iron ion metabolic Network.**

**Supplementary information of sever implementation**

*Frontend.* The website (http://scflux.org) is implemented in Python using the Django framework, and it uses the SQLite (https://www.sqlite.org/) database for persistent storage and retrieval of requested data. The Nginx (http://nginx.org/) web server acts as a secure application gateway. It acts as a load balancer, a reverse proxy, and a caching mechanism. It handles simultaneous connections and accelerates the loading of static files such as images. In addition to serving static content and encrypt requests, it also balance the load of a large number of requests and distribute those requests evenly across many uWSGI (https://uwsgi-docs.readthedocs.io/) instances. Then uWSGI translates requests from Nginx into a format that the Python application understands. The Nginx + uWSGI architecture enables the application to handle a large number of simultaneous connections.

The user interface of the website is designed using the Bootstrap framework, the jQuery JavaScript library, and their extensions. Icons are taken from Font Awesome (https://fontawesome.com). And the code highlighting of web pages is implemented based on prism.js (https://prismjs.com/).

*Backend. Input data preprocessing* is based on Pandas (https://pandas.pydata.org/) and NumPy (https://numpy.org/). It mainly includes illegal character detection, reformatting and normalization of input data. If the input data does not meet the analysis requirements, a warning message will be given on a case-by-case basis. The graph neural network model is implemented based on PyTorch (https://pytorch.org/), Pandas and NumPy. The loss curve is drawn using Matplotlib (https://matplotlib.org/). If the data is sparse, an imputation procedure is recommended, the default imputation method is MAGIC. Data processing procedure composes three major steps: (1) an evaluation step that checks if the input is in a correct format. Warnings will be returned if the data format does not fit the requirement of scFLUX; (2) data normalization and imputation by using MAGIC; and (3) pass the input data and user selected metabolic network to the flux estimator.

*The Flux estimation procedure*. The graph neural network model is implemented by using PyTorch (https://pytorch.org/), Pandas and NumPy. The factor graph model of each metabolic network was built, in which each module is a variable, and each intermediate metabolite is a factor. Parallel small three-layer neural networks were first constructed to model the non-linear dependency between gene expression involved in and flux of modules. Noted, one neural network is constructed for each module. The loss function is further constructed based on the in-/out-flux of each intermediate metabolites and minimized by using the Adam method, which is the most efficient stochastic optimization approach in the built-in optimizer of

Pytorch. The curve of the loss though the training process is drawn by using Matplotlib (https://matplotlib.org/) and shown on the result page.

*Job management.* Analysis tasks can run asynchronously, and the management of these tasks is based on the Celery (https://docs.celeryproject.org/), a distributed task queue. While Redis (https://redis.io/) is used to store messages for Celery workers, and store results coming off the Celery queues. Based on this mechanism, users can run multiple analysis tasks simultaneously. When a user submits a task, a unique task ID and link will be generated, by which the user can track the progress percentage of the analysis task, and when the task is completed, the user can access, download, or share the analysis results. Analysis results will be stored on the server for at least two weeks if storage space permits. The list of user's tasks is implemented by using a cookie mechanism. Therefore, it only lists previous tasks submitted by the user using the same browser. However, access via link has no such restrictions.

**Background of the scFEA method**

We have recently developed a computational method to estimate cell-wise metabolic fluxome (flux distribution of the whole metabolic network) by using single-cell RNA-seq data (scRNA-seq) [3] (*Genome Research* 2021, highlighted by *RECOMB*, *ICIBM*, and *ENAR*). To the best of our knowledge, this is the first and only method can estimate sample-wise flux distribution of a metabolic network in single cell resolution. scFEA utilizes a factor graph base representation of metabolic network and a novel graph neural network (GNN) model for flux estimation, by assuming (1) metabolic flux can be modeled as a neural network of the genes involved in neighboring reactions, and (2) minimization of the flux imbalance of intermediate metabolites. Compared to existing methods, scFEA is the only method designed for disease study, which specifically (1) models the nonlinear dependency between gene expression and metabolic flux, (2) assess flux in each single cell, and (3) enable certain imbalance of intermediate metabolites in highly intertwined disease microenvironment, as detailed in the table below. We also developed functions to compute (i) accumulation or depletion of metabolites, (ii) the impact of each gene on metabolic fluxome, and (iii) the subset of cells having distinct variation of certain metabolic modules [3]. In the last six months, the software package of scFEA received more than 50 inquires of usage or collaborations and more than 500 downloads [4].

| Comparison of scFEA vs other metabolic analysis models | *Elementary mode (EM)* | *Flux Balance Analysis (FBA)* | *Single sample pathway analysis* | *scFEA* |
|---|---|---|---|---|
| **Loss Assumption** | stringent flux balance | stringent flux balance | average of gene expression | **quadratic loss of flux balance** |
| **Model** | topology | dynamic programming | pathway enrichment score | **mass carrying GNN** |
| **Network representation** | elementary mode | simple graph | gene set, pathway | **factor graph** |
| **#Variables** | $>10^5$ modes | $\sim 10 - 10^3$ reactions | $\sim 10^2 - 10^3$ pathways | **$\sim 10 - 10^3$ modules** |
| **Representative Methods** | elementary mode | COMPASS, scFBA | ssGSEA | **scFEA** |
| **Targeted pathways** | whole network | partial network | predefined pathways | **whole or any sub-network** |
| **Resolution** | bulk tissue | cell cluster | single cell | **single cell** |
| **Approximate kinetic model** | No | No | No | **Yes** |
| **Disease specific metabolism** | No | No | No | **Yes** |
| **Predict metabolomic change** | No | No | No | **Yes** |

**Previous validations of the scFEA method**

As scFEA is the first attempt of using graph neural network and a quadratic loss function for a sample-wise metabolic flux estimation. To validate the method assumptions and optimize loss forms, we established an experiment platform by collecting matched scRNA-seq and tissue metabolomics data of Pa03c, a patient derived pancreatic cancer cell line under perturbed oxygen, oxidative stress and knock-down of metabolic genes [5]. scFEA and their functions has been validated by qRT-PCR and knock-down experiments on this system [3].

Specifically, we observed a Pearson correlation coefficient (PCC) of 0.86 (p=0.006) between scFEA predicted and experimentally observed flux change [5]. On the other hand, we also correlated the observed metabolomic change with the averaged expression change of the enzymes catalyzing each reaction. No significant correlation was observed (PCC=-0.03, p=0.943), suggesting that single cell gene expression alone doesn't produce a good estimate of single cell metabolomic landscape. In addition, we also examined the ssGSEA (single sample gene set enrichment analysis) by using the reduced metabolic modules as gene sets [6]. Averaged ssGSEA score considered as the activity the metabolic modules estimated by ssGSEA. A correlation 0.42 (p=0.299) was observed between the difference in averaged ssGSEA score vs the metabolomics change. We speculate ssGSEA does not work very well on estimate single cell metabolic flux because the weight of each gene in one module was fixed by their expression level and no flux balance condition was utilized.

**Hyperparameter of scFEA**

We and a few users have tested the robustness of the hyperparameters of scFEA. We empirically found that setting the hyperparameters as $\alpha = 1$, $\beta = 0.1$, $\gamma = 1$ achieved (1) low total loss, (2) good biological expandability, (3) high goodness of fitting of the consistency between gene expression and predicted flux, and (4) fast convergency of the model training, on TCGA and CCLE bulk tissue RNA-seq data and more than 20 scRNA-seq data. In scFLUX, to avoid further confusion and provide a simple analysis environment, we fixed this hyperparameter. Users can adjust hyperparameters by using the standalone package of scFLUX.

**Public scRNA-seq data used in this study**

Human cancer scRNA-seq Data

GSE72056: This dataset is collected on human melanoma tissues. The original paper provided cell classification and annotations including B cells, cancer-associated fibroblast (CAF) cells, endothelial cells, macrophage cells, malignant cells, NK cells, T cells, and unknown cells.

GSE103322: This dataset is collected on head and neck cancer tissues. The original paper provided cell classification and annotations including B cells, dendritic cells, endothelial cells, fibroblast cells, macrophage cells, malignant cells, mast cells, myocyte cells, and T cells. Notably, as indicated by the original work, malignant cells have high intertumoral heterogeneity.

GSE115978: This dataset is collected on 31 human melanoma tumors. Isolated immune and non-immune cells by FACS based on CD45 staining for T cell exclusion associated target.

GSE140182: This dataset is collected on 5 advanced human gastric cancer sample. It includes 4 malignant ascites and 1 cerebrospinal fluid metastasis. After data quality control, total 162 cells from AGC patients, 97 cells from M1, and 45 cells from M2 were used.

GSE144735: This dataset is collected on 6 Belgian colorectal cancer patients (CRC). After quality control, 27,414, cells were used.

GSE145137: This dataset is collected on chemotherapy-resistant muscle-invasive urothelial bladder cancer. scRNA sequencing data were acquired from a patient and patient-derived xenografts. Total 3,934 cell pass the quality control.

GSE146409: This dataset is collected on liver tumor. Six patients who underwent liver resection (colorectal metastasis, primary cholangiocarcinoma, and benign liver cyst). scRNA sequencing using MARSseq.

GSE151530: This dataset is collected on liver cancer and a total of 46 tumor samples were profiled.

Mouse injected tumor scRNA-seq

GSE132582: This dataset is collected on pancreatic cancer. scRNA sequencing were performed on a total of three samples.

GSE136206: This dataset is collected on triple negative breast cancer. Total four samples were collected with detailed ID: T11-Apobec-7daytreated, T11-Apobec-Nottreated, KPB25Luv-7daytreated, KPB25Luv-Nottreated.

ROSMAP data

This dataset includes snRNA-seq data collected from 24 Alzheimer's disease (AD) patient and 24 healthy donor brain samples, which was generated from the Religious Orders Study (ROS) or the Rush Memory and Aging Project (MAP), mainly focus on the Alzheimer's disease research. The dataset was download from RADC Research Resource Sharing Hub (https://www.radc.rush.edu/)

**Detailed analysis and results of the case studies.**

*Cancer Microenvironment scRNA-seq data*

We computed metabolic flux of the GlucoseGlutamineOpen network by using the 8 human and 3 mouse cancer scRNA-seq data. In each data set, we examined the flux of glucose metabolism into TCA cycle, lactate, nucleic acids and serine, which are shown in **Supplementary Figure S8A and S8B**. We also estimated the cell proliferation rate by using ssGSEA against the cell proliferation GO term and computed the correlation between cell proliferation rate and glucose consumption rate in each data set (**Supplementary Figure S8C**).

*ROSMAP Alzheimer's Disease snRNA-seq data*

We computed metabolic flux of the M171 network by using the ROSMAP snRNAs-seq data. We identified the metabolic activity of neuron cells is much higher than other cell types. We further computed the averaged concentration (metabolomic) change of all intermediate metabolites between the neuron cells from AD patients and healthy donors. **Supplementary Table S2** listed the top varied metabolites in AD neuron cells vs healthy donor controls.

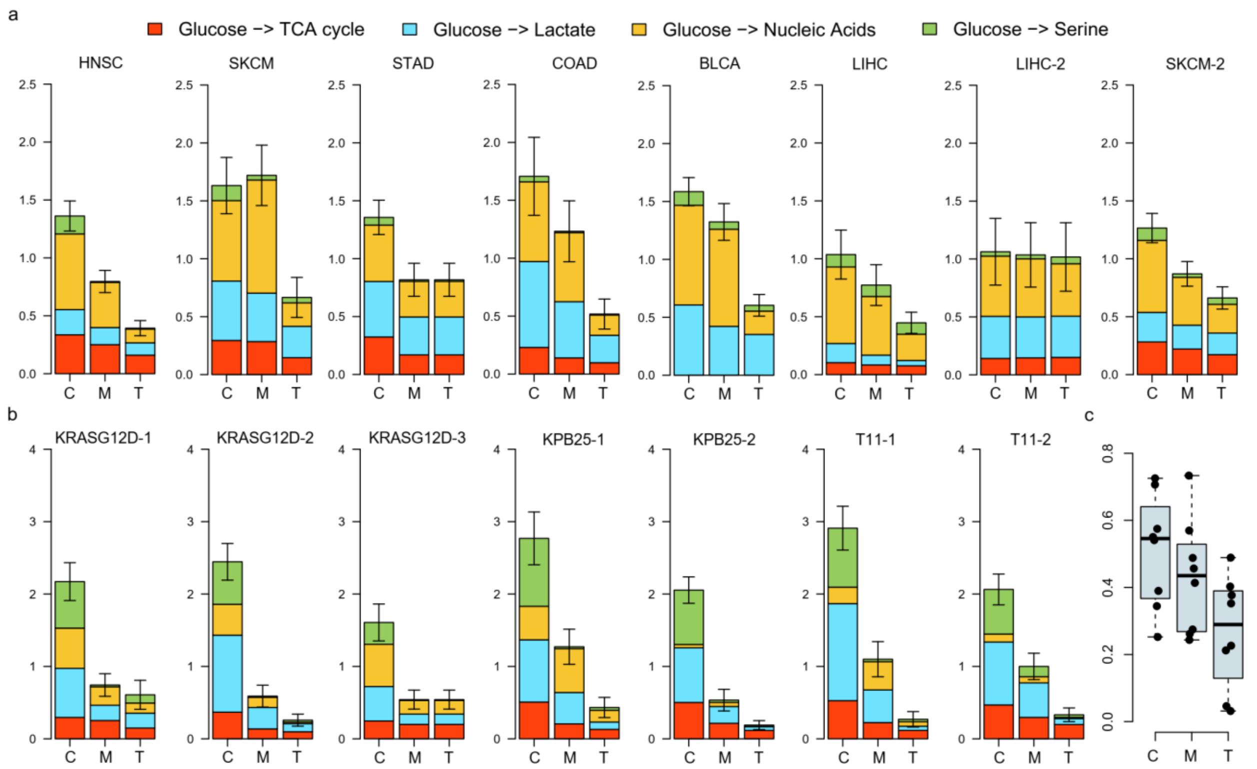


**Supplementary Figure S8. Case study on scRNA-seq data of cancer microenvironment.** Predicted flux of glucose metabolisms in cancer (C), myeloid (M) and T (T) cells in (A) eight types of human cancer tissues and (B) orthotopic mouse tumors, where the height of each colored segment represents the level of the flux into a specific branch. (C) Correlations between glucose consumption and cell proliferation rates in human cancer, myeloid and T cells based on scRNA-seq data.

| Metabolites | Abundance | p.value | Formula |
|---|---|---|---|
| **Succinyl-CoA** | 0.707 | <1e-300 | $C_{25}H_{40}N_7O_{19}P_3S$ |
| **Citrate** | 0.645 | <1e-300 | $C_6H_8O_7$ |
| **Fumarate** | 0.381 | <1e-300 | $C_4H_4O_4$ |
| **Succinate** | 0.304 | 1.4e-203 | $C_4H_6O_4$ |
| **Malate** | 0.282 | 1.1e-217 | $C_4H_6O_5$ |
| **Glutamine** | 0.255 | 1.5e-83 | $C_5H_{10}N_2O_3$ |
| **GABA** | 0.181 | 1.6e-66 | $C_4H_9NO_2$ |
| **Methionine** | -0.168 | 4.3e-58 | $C_5H_{11}NO_2S$ |
| **Serine** | -0.192 | 4.5e-93 | $C_3H_7NO_3$ |
| **Leucine** | -0.194 | 7.4e-58 | $C_6H_{13}NO_2$ |
| **CDP** | -0.313 | 1.3e-27 | $C_9H_{15}N_3O_{11}P_2$ |
| **Glucose** | -0.552 | 9.04e-63 | $C_6H_{12}O_6$ |
| **G6P** | -0.732 | 1.2e-274 | $C_6H_{13}O_9P$ |
| **Aspartate** | -0.853 | <1e-300 | $C_4H_7NO_4$ |

**Supplementary Table S2. Metabolites that have top abundance change in AD brain vs normal brain predicted by using ROSMAP snRNA-seq data.**
